# Aspirin-mediated DKK-1 increase rescues Wnt-driven stem-like phenotype in human intestinal organoids

**DOI:** 10.1101/809517

**Authors:** Karen Dunbar, Asta Valanciute, Vidya Rajasekaran, Thomas Jamieson, Paz Freile Vinuela, Ana Lima, James Blackmur, Anna-Maria Ochocka-Fox, Mark J. Arends, Owen J. Sansom, Kevin B Myant, Susan M. Farrington, Malcolm G. Dunlop, Farhat V.N. Din

## Abstract

Aspirin reduces the incidence and mortality of colorectal cancer (CRC). Wnt signalling drives CRC development from initiation to progression through regulation of epithelial-mesenchymal transition (EMT) and cancer stem cell populations (CSC). Here, we investigated whether aspirin can rescue these pro-invasive phenotypes associated with CRC progression in Wnt-driven human and mouse intestinal organoids. Aspirin rescues the Wnt-driven cystic organoid phenotype by promoting budding in mouse and human Apc deficient organoids, which is paralleled by decreased stem cell marker expression. Aspirin-mediated Wnt inhibition in Apc^Min/+^ mice is associated with EMT inhibition and decreased cell migration, invasion and motility in CRC cell lines. Chemical Wnt activation induces EMT and stem-like alterations in CRC cells, which are rescued by aspirin. Aspirin increases expression of the Wnt antagonist Dickkopf-1 (DKK-1) in CRC cells and organoids derived from FAP patients. We provide evidence of phenotypic biomarkers of aspirin response with an increased epithelial and reduced stem-like state mediated by an increase in DKK-1. Thus we highlight a novel mechanism of aspirin-mediated Wnt inhibition and potential phenotypic and molecular biomarkers for trials.

## INTRODUCTION

Colorectal cancer (CRC) is the 4^th^ leading cause of cancer death world-wide (Ferley et al.). This highlights the need to understand and counteract the molecular pathways underlying CRC initiation and progression. Compelling data show that aspirin decreases CRC incidence by 40-50% (Din et al., 2010; Rothwell et al., 2010) and that post-diagnosis intake reduces mortality suggesting delayed disease progression (Bastiaannet et al., 2012; Chia et al., 2012). Understanding the biology responsible for the protective effect is key to developing biomarker-led approaches for its rational clinical use.

Canonical Wnt signaling plays a critical role in cell survival, stem cell and gastrointestinal homeostasis (Logan and Nusse, 2004). Wnt signaling is activated and inhibited by numerous extracellular ligands such as the Wnts and Dickkopf proteins respectively (Cruciat and Niehrs, 2013). Activation by Wnt ligands protects cytoplasmic β-catenin from destruction, leading to its nuclear translocation and TCF/LEF-dependent Wnt target gene induction (Logan and Nusse, 2004). The Wnt inhibitor Dickkopf-1 (DKK-1) is downregulated, during the adenoma-carcinoma sequence with low expression correlating with increased tumorigenesis and angiogenesis (Liu et al., 2015). The importance of dysregulated Wnt in CRC initiation, driven by Apc mutations and less frequently □-catenin in sporadic CRC, is well-established in sporadic CRC and familial adenomatous polyposis (FAP). Growing evidence suggests Wnt signaling regulates mechanisms associated with CRC progression such as epithelial-mesenchymal transition (EMT) and cancer stem cells (CSC) (Cai et al., 2018).

Malignant cells co-opt EMT regulatory machinery to gain motility and, ultimately, metastatic properties (Kalluri and Neilson, 2003). There is progressive loss of epithelial features with loss of cell-cell junctions and cell polarity, and replacement by an invasive mesenchymal phenotype (Kalluri and Neilson, 2003). The plasticity of this process leads to partial phenotypes and mesenchymal-epithelial transition which promotes metastases to establish (Dawson and Lugli, 2015). Poor CRC survival and metastases are associated with EMT-inducing gene expression and mesenchymal histology (Spaderna et al., 2006; Ueno et al., 2014). Furthermore, EMT induction by the transcription factors Snail, Slug or Twist triggers the acquirement of a stem-like phenotype in several cancers (Mani et al., 2008). Cancer stem cells (CSCs), a subpopulation of self-renewing cells, exist in numerous cancers including colorectal (Ricci-Vitiani et al., 2007). CSCs may arise from cancer cell dedifferentiation or from normal colonic stem cell transformation through increased intrinsic tumorigenesis or in response to the microenvironment (Mani et al., 2008). Importantly diverse pathways dysregulated in CRC, including Wnt signaling, converge to regulate both EMT and CSCs highlighting pro-malignant phenotypes that may be monitored for drug response. There is an accelerated drive towards disease-relevant phenotypic screening, rather than the limited clinical success of target-based approaches for drug discovery (Jones and Bunnage, 2017).

Despite the importance of Wnt signaling in CRC, there are no specific inhibitors used clinically (Tai et al., 2015). Wnt inhibitor development has been challenging due to the complex and temporal role of Wnt in development and homeostasis, and the presence of cross-talk from other Wnt-regulating pathways. Hence, a drug such as aspirin which either targets multiple nodes within Wnt signaling or multiple pathways to combat redundancy may be more effective (Gala and Chan, 2015). The efficacy of aspirin is likely due to its ability to modulate multiple aberrant CRC networks including mTOR and Wnt signaling (Bos et al., 2006; Din et al., 2012). Whilst aspirin reduces nuclear β-catenin expression to inhibit canonical Wnt signaling in CRC cell lines (Bos et al., 2006), the role of aspirin on EMT and CSC phenotypes due to this inhibition has not been determined. Here we investigate the inhibitory effect of aspirin on metastases-promoting mechanisms in CRC and whether these effects can be attributed to inhibition of canonical Wnt signalling.

## RESULTS

### Aspirin reverses the Wnt-driven cystic organoid phenotype

Intestinal organoids are an invaluable model for studying intestinal stem cells as the familiar budding projections represent crypt-like structures, initiated by the presence of an Lgr5^+^ stem cell and a microenvironment generated by growth factors (Sato et al., 2011). Local Wnt signaling, critical for normal stem cell function and bud formation, is dysregulated in Apc^Min/+^, Apc^flox/flox^ and FAP adenoma organoids which grow in a characteristic cystic manner without budding structures (Sup. Fig. 1A) (Sato et al., 2011). Given clinical trials of aspirin in sporadic adenoma and familial syndromes show benefit we investigated whether aspirin alters the Wnt-driven phenotype in organoids. Treating small intestinal organoids grown from an Apc^flox/flox^ mouse with 2mM aspirin for 12 days increased the percentage of non-cystic organoids, identified as those organoids containing buds, compared to the untreated population (Fig. 1A and Video 1a, b). These effects were translated to human organoids grown from non-adenomatous (macroscopically normal) and adenomatous colonic tissue from FAP patients. Treatment with 0.5mM aspirin for 29 days increased the percentage of budding organoids derived from both non-adenomatous and adenomatous colonic mucosa (Fig. 1B). Additionally, treatment of Apc^Min/+^ mice with aspirin for 4 weeks *in vivo* produced organoids that grew with a budding phenotype compared to untreated Apc^Min/+^ intestinal organoids (Sup. Fig. 1B). Hence, we present robust evidence that aspirin rescues the cystic phenotype characteristic of constitutively active Wnt signaling due to a mutant Apc background.

**Figure 1:**
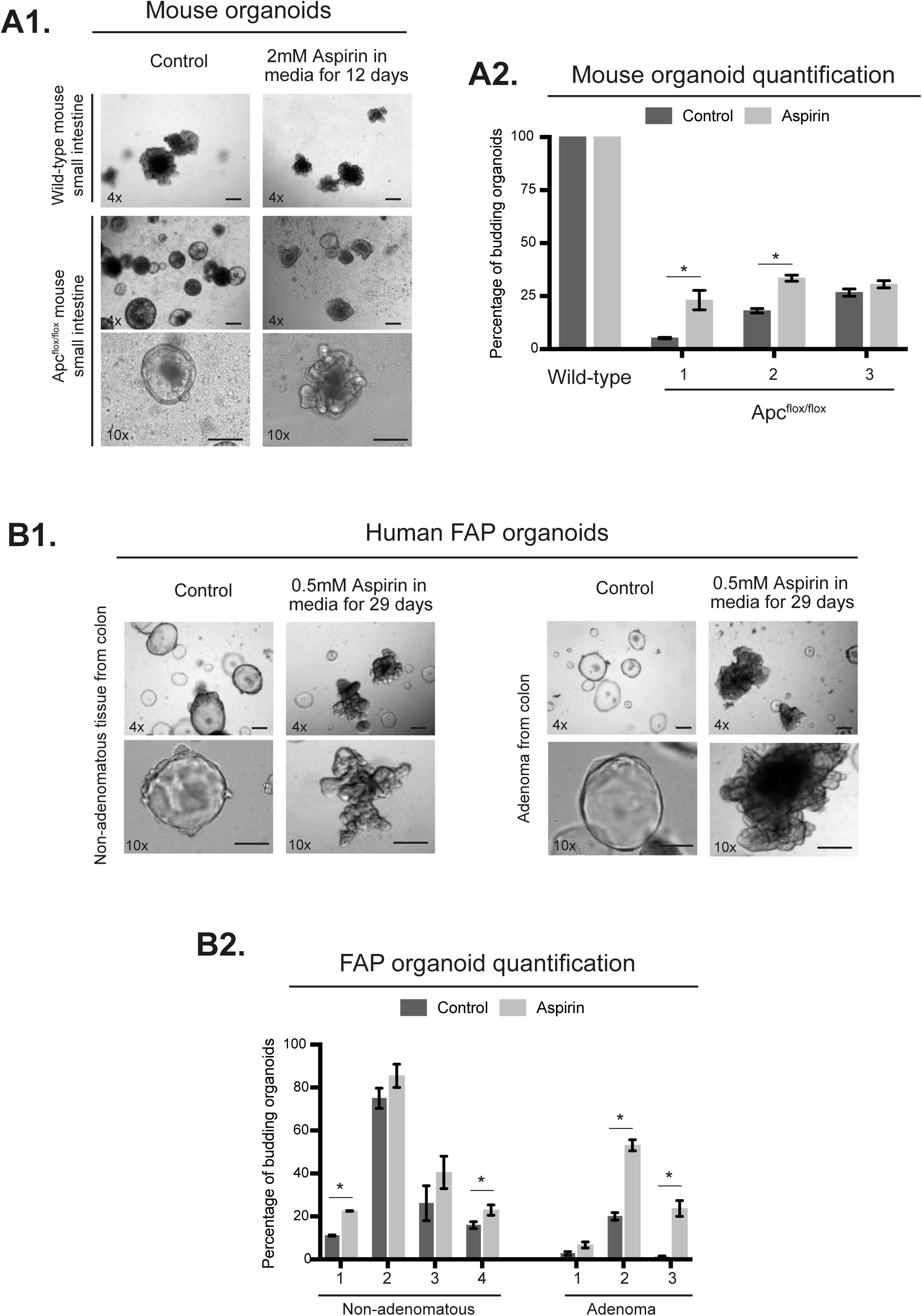
Aspirin alters the organoid budding phenotype. Brightfield images of small intestinal organoids from Wild-type (C57BL/6j) and Apc^flox/flox^ mouse organoids treated with 2mM aspirin for 12 days (A1). Quantification of the percentage of non-cystic organoids from one wild-type (C57BL/6j) mouse and three Apc^flox/flox^ mice (A2). Brightfield images of organoids from human FAP non-adenomatous and adenomatous colon tissue treated with 0.5mM aspirin for 29 days (B1) and quantification of the percentage of budding organoids (B2). Quantification of four biological replicates from non-adenomatous derived organoids and three biological replicates from adenoma derived organoids. Microscope objective magnification noted in bottom left of image. Error bars represent standard error of mean. Significance determined by unpaired students t-test. * p-value <0.05. Scale bar = 200um. FAP: Familial adenomatous polyposis. NA: Non-adenomatous.

### Aspirin decreases stem cell marker expression and reduces Wnt signaling

The phenotype changes in these organoid models suggest aspirin-mediated effects on either the Lgr5^+^ stem cell population or the Wnt signaling gradients required for efficient organoid budding (Sato and Clevers, 2013). *Lgr5*, a Wnt target gene and receptor for the Wnt agonist R-spondin, is the principle stem cell marker in intestinal tissue but other proteins, such as *TROY*, have also been validated as intestinal stem cell makers (Barker et al., 2007). Basal *Lgr5* and *TROY* expression is increased in Apc^flox/flox^ organoids compared to wild-type organoids, illustrating the increased number of stem cells in Apc mutant tissue and aspirin treatment reduces transcript expression of both markers in Apc^flox/flox^ organoids (Supp. Fig. 1C). Utilising RNAscope technology we established the relative expression of *Lgr5* and *TROY* RNA transcripts per organoid with aspirin treatment reducing expression of both markers (Fig. 2A). In the 4-week treatment Apc^Min/+^ model, crypts isolated from aspirin-treated mice also showed a reduction in *Lgr5* and *TROY* transcript expression compared to untreated mice (Sup. Fig. 1D). Aspirin treatment significantly reduced *Lgr5* transcript expression in colonic organoids derived from human patients with varying pathology including; normal mucosa, sporadic CRC and FAP CRC (Fig. 2B). In FAP human organoids from non-adenomatous and adenomatous tissue Lgr5 protein expression was also reduced with aspirin (Fig. 2C). These findings were replicated *in vivo*, with the treatment of Apc^Min/+^ mice with aspirin for 4 weeks decreasing the number of *Lgr5* RNA transcripts per adenoma (Fig. 2D). The aspirin-mediated decrease in stem cell markers suggests alterations in the stem cell niche, specifically the Paneth cells which are adjacent to Lgr5^+^ cells. Paneth cell loss drives a reduction in Lgr5^+^ stem cells highlighting their requirement for stem cell homeostasis (Farin et al., 2012). In Apc^Min/+^ mice treated for 7 days, aspirin decreased the number of Paneth cells in small intestinal adenomas, as identified by lysozyme immunohistochemistry (Fig. 2E).

**Figure 2:**
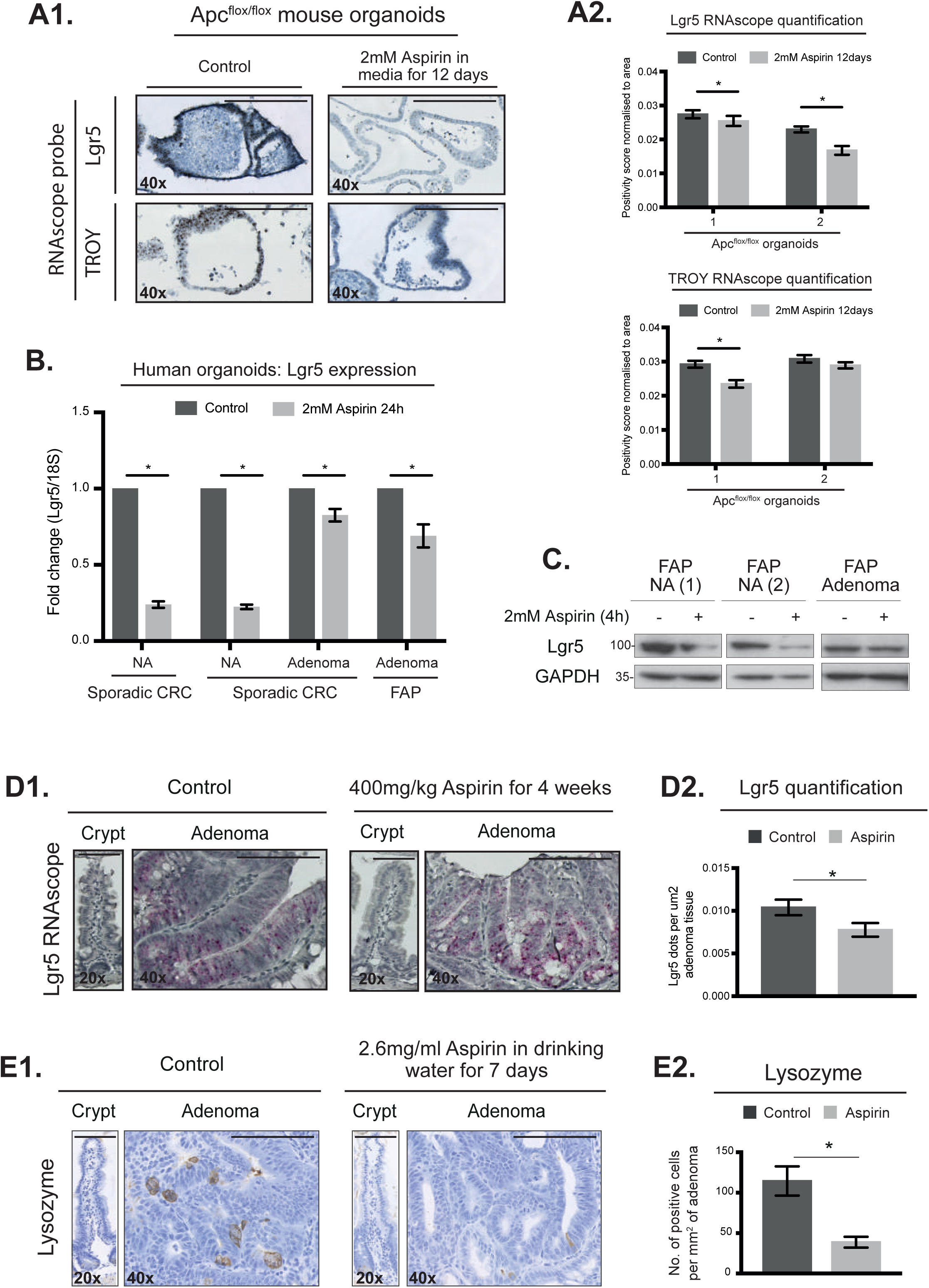
Aspirin reduces stem cell marker expression in organoids and *in vivo*. Images of Apc^flox/flox^ mouse organoids treated with 2mM aspirin for 12 days and stained with RNAscope probes for either *Lgr5* or *TROY* (A1). Quantification from 2 biological replicates (between 25-90 organoids per condition) and presented as positivity score normalised to area (A2). RNA transcript expression of *Lgr5* in organoids from human colonic tissue (two sporadic CRC patients and one FAP patient) treated with 2mM Aspirin 24hours (B). Immunoblotting of Lgr5 protein in human colonic organoids, three patient samples, treated with 2mM aspirin for 4 hours (C). RNAscope staining images (D1) and quantification (D2) of control and aspirin treated (400mg/kg aspirin for 4 weeks) Apc^Min/+^ mouse tissue for *Lgr5*. Quantification of the number of *Lgr5* dots per um^2^ of adenoma tissue (24 control and 30 aspirin treated adenomas) from cohort of five Apc^Min/+^ control and four Apc^Min/+^ aspirin treated mice. Immunohistochemistry images (E1) and quantification (E2) of control and aspirin treated (2.6mg/ml aspirin in drinking water for 7 days) Apc^Min/+^ mouse tissue for Lysozyme. Immunohistochemistry quantification for 50 control adenomas from five Apc^Min/+^ mice and for 34 aspirin treated adenomas from four Apc^Min/+^ mice. Microscope objective magnification noted in bottom left of image. Error bars represent standard error of mean. Significance determined by unpaired students t-test. * p-value <0.05. Scale bar = 50um. FAP: Familial adenomatous polyposis. NA: Non-adenomatous.

The predominant pathway responsible for stem cell niche maintenance is Wnt signaling which is known to be inhibited by aspirin (Sato and Clevers, 2013). We confirmed this by demonstrating aspirin decreased β-catenin protein expression in small intestinal adenomas from Apc^Min/+^ mice (Fig. 3A). Aspirin treatment also reduced transcript expression of *Tcf7*, the transcriptional co-activator of β-catenin (Clevers and Nusse, 2012), in Apc^Min/+^ crypts and Apc^flox/flox^ organoids (Sup. Fig. 1D). Wnt signaling has been implicated in the regulation of metastatic mechanisms including EMT. It is noteworthy that whilst Apc^Min/+^ adenomas are predominately epithelial in nature, we observed a small number of vimentin positive cells within these epithelial populations suggesting cells undergoing EMT (Sup. Fig. 2). We observed changes in the expression of E-cadherin and Vimentin in small intestinal adenomas from Apc^Min/+^ mice with aspirin increasing E-cadherin and reducing Vimentin expression (Fig. 3A). These observations indicate that aspirin promotes a more epithelial phenotype *in vivo* and may contribute to inhibition of EMT progression through aspirin-mediated Wnt inhibition.

**Figure 3:**
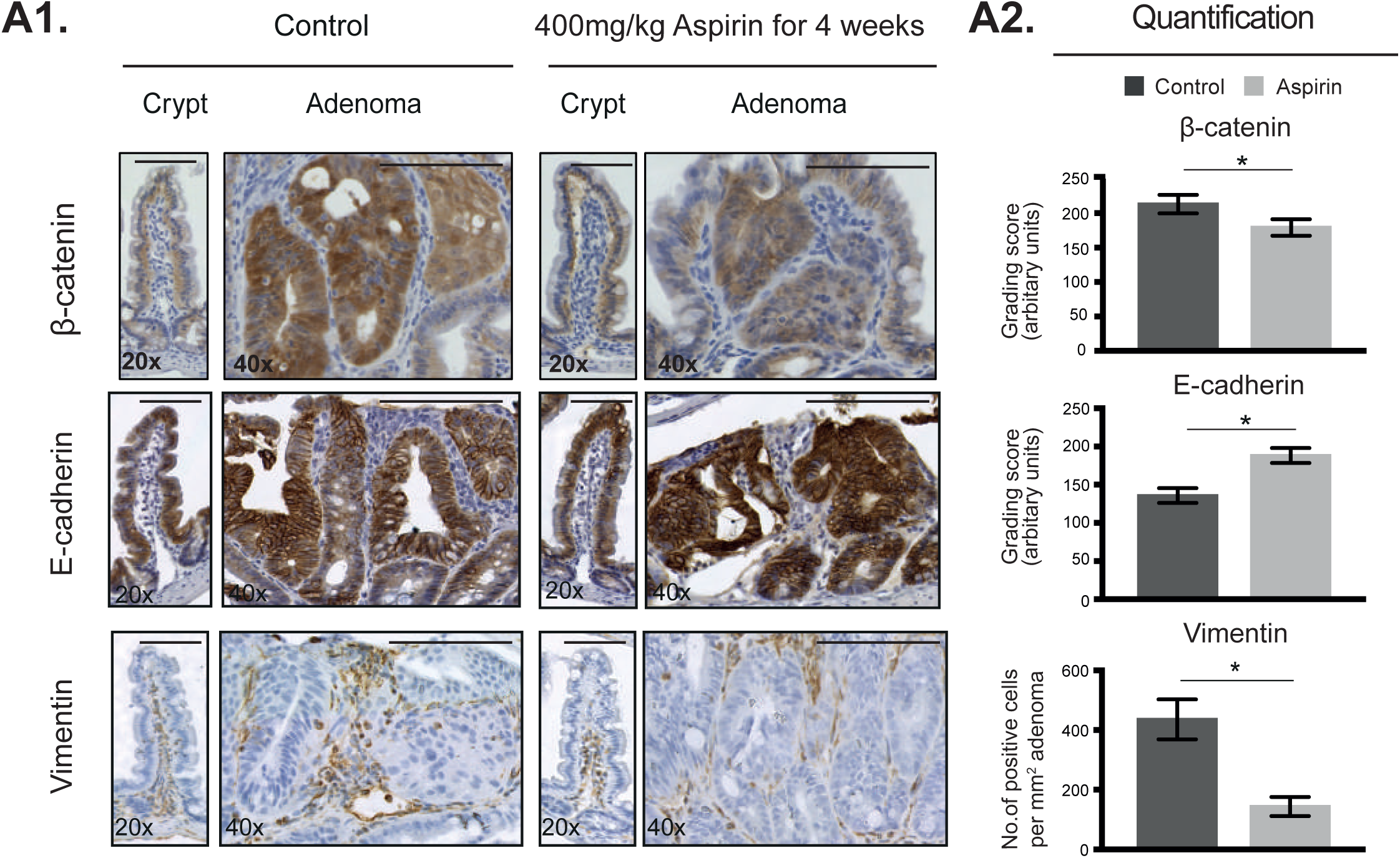
Aspirin reduces β-catenin and EMT markers *in vivo*. Immunohistochemistry images (A1) and quantification (A2) of control and aspirin treated (400mg/kg aspirin for 4 weeks) Apc^Min/+^ mouse tissue for β-catenin, E-cadherin and Vimentin. Immunohistochemistry quantification of β-catenin (21 control and 30 aspirin treated adenomas), E-cadherin (37 control and 23 aspirin treated adenomas) and Vimentin (29 control and 16 aspirin treated adenomas) from cohort of five Apc^Min/+^ control and four Apc^Min/+^ aspirin treated mice. Microscope objective magnification noted in bottom left of image. Error bars represent standard error of mean. Significance determined by unpaired students t-test. * p-value <0.05. Scale bar = 50um.

### Aspirin inhibits Wnt and EMT whilst reducing migration and invasion of CRC cells

These *in vivo* observations were confirmed in Colo205 cells with aspirin reducing expression of both β-catenin and its targets; c-myc and Lgr5 whilst increasing E-cadherin expression (Fig. 4A). Aspirin treatment increased E-cadherin protein and transcript expression in HCT116 and Colo205 cells (Sup. Fig. 3A, B). An additional epithelial marker, zona-occludens 1 (ZO-1), was also increased upon aspirin treatment (Sup. Fig. 3C). Aspirin exposure reduced expression of Snail, a key transcription factor which represses E-cadherin to drive EMT, in parallel to increasing cytoplasmic E-cadherin expression in HCT116 cells (Sup. Fig. 3D). These results confirm that aspirin is promoting an enhanced epithelial phenotype.

**Figure 4:**
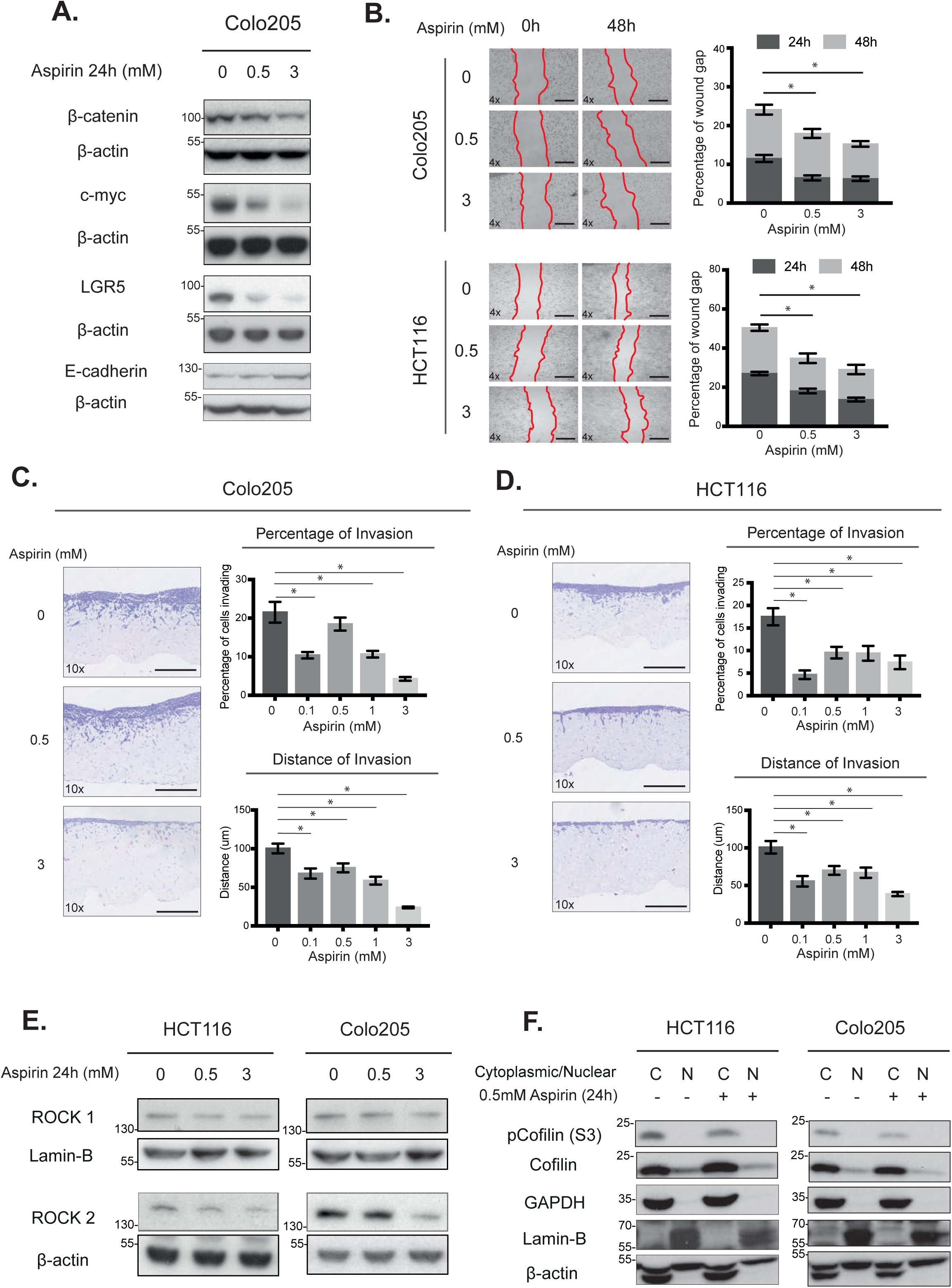
Aspirin reduces Wnt signaling, migration and invasion in CRC cells. Immunoblotting in Colo205 cells treated with 0.5 or 3mM aspirin for 24 hours (A). Brightfield images and quantification of wound closure assays over 48 hours in 0.5% serum in Colo205 and HCT116 cells (B). Red outline indicates wound edges. Scale bar = 100um. Wound closure assay data representative of three biological replicates each containing three technical replicates. Brightfield images and quantification of the percentage of cells invading and the maximum distance invaded in organotypic invasion assays over 7 days with Colo205 (C) and HCT116 (D) cells. Scale bar = 200um. Invasion assay data representative of three biological replicates each containing two technical replicates. Immunoblotting of HCT116 and Colo205 cells after 24hour treatment with 0.5 or 3mM aspirin (E). Immunoblotting of cytoplasmic/nuclear extracts treated with 0.5mM aspirin for 24 hours in HCT116 and Colo205 cells (F). Immunoblotting data representative of three biological replicates. Microscope objective magnification noted in bottom left of image. Error bars represent standard error of mean. Significance determined by unpaired students t-test. * p-value <0.05

Increased migratory and invasive capabilities are characteristic traits of cells undergoing EMT and disease progression. Aspirin reduced wound closure in HCT116 and Colo205 cells grown in both low (0.5%) and normal (10%) serum conditions (Fig. 4B and Sup. Fig. 4A, B). In low serum conditions cellular proliferation is inhibited, hence effects on wound closure more closely reflect alterations in migratory capacity rather than proliferation thus giving increased confidence that aspirin inhibits CRC cell migration. This is particularly important given aspirin has known anti-proliferative effects especially at higher concentrations (Sup. Fig. 5). Cellular invasion was modelled using organotypic invasion assays in which cells invade collagen-fibroblast matrices. Aspirin decreased both the distance invaded and the overall percentage of HCT116 and Colo205 cells invading, which is normalised to the non-invading population to remove any anti-proliferative bias (Fig. 4C, D).

**Figure 5:**
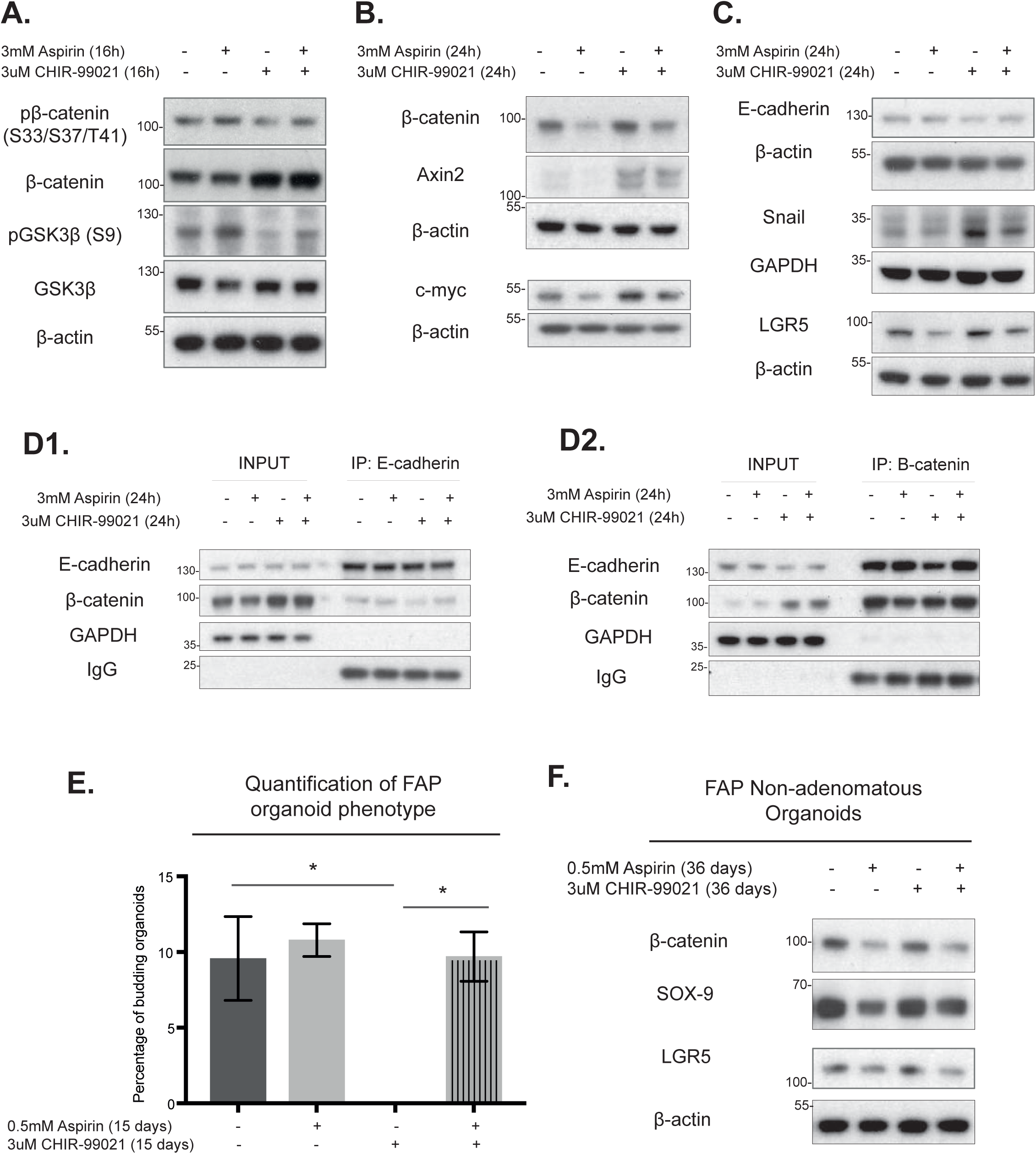
Aspirin reduces Wnt driven mesenchymal and stem marker expression in CRC cell lines and FAP organoids. Immunoblotting of phospho-β-catenin (S33/S37/T41), β-catenin, phospho-GSK3β (S9) and GSK3β in HCT116 cells treated with 3mM aspirin, 3μM CHIR-99021 or combination for 16 hours (A). Immunoblotting of β-catenin, Axin2 and c-myc with 3mM aspirin, 3μM CHIR-99021 or combination for 24 hours in HCT116 cells (B). Immunoblotting of E-cadherin, Snail and LGR5 with 3mM aspirin, 3μM CHIR-99021 or combination for 24 hours in HCT116 cells (C). Immunoprecipitation using E-cadherin antibody in HCT116 cells treated with either 3mM aspirin, 3μM CHIR-99021 or combination for 24 hours (D1). Immunoprecipitation using β-catenin antibody in HCT116 cells treated with either 3mM aspirin, 3μM CHIR-99021 or combination for 24 hours (D2). Immunoblotting data representative of three biological replicates. Quantification of the percentage of budding organoids derived from human FAP patient colon mucosa and treated with 0.5mM aspirin, 3μM CHIR-99021 or combination for 15 days in vitro (E). Immunoblotting of colonic organoids derived from single human FAP patient and treated with 0.5mM aspirin, 3μM CHIR-99021 or combination for 36 days in vitro (F). Error bars represent standard error of mean. Significance determined by unpaired students t-test. * p-value <0.05.

These effects translated to cellular motility with aspirin reducing the distance single CRC cells travelled in 24 hours (Sup. Fig. 4C, D). RhoA-Rac1 signaling has been specifically implicated in regulating motility in CRC cells undergoing EMT (Gulhati et al., 2011). We focussed on the downstream effectors of the RhoA-Rac1 pathway; Rho-associated kinases 1 and 2 (ROCK1 and 2), cofilin and F-actin (Rath and Olson, 2012). Aspirin reduced the protein expression of ROCK1 and ROCK2, (Fig. 4E) reduced levels of phosphorylated cofilin in CRC cells (Fig. 4F) and reduced F-actin expression (Sup. Fig. 4E). Modulation of the balance between phosphorylated and unphosphorylated cofilin alters F-actin stabilisation and cycling which is required for cell movement (Scott et al., 2010). These results demonstrate that aspirin can modulate several cell traits commonly associated with EMT and disease progression in CRC cell lines.

### Aspirin treatment rescues Wnt-driven EMT and stem cell changes in CRC cells

To investigate whether aspirin could specifically reverse a Wnt-driven EMT and associated stem cell alterations we utilised the GSK-3β inhibitor, CHIR-99021 to hyper-activate Wnt signaling. Aspirin treatment abrogates CHIR-99021-mediated Wnt activation by increasing GSK-3β and β-catenin phosphorylation (Fig. 5A). The CHIR-99021 mediated increase in β-catenin and its target gene expression, Axin2 and c-myc, is reversed upon aspirin exposure (Fig. 5B). Wnt activation promotes a mesenchymal stem-like phenotype with decreased E-cadherin and increased Snail and Lgr5 expression, which was attenuated by aspirin in HCT116 cells (Fig. 5C). Increased E-cadherin has been shown to buffer excessive β-catenin thus limiting hyper-activated Wnt and further promote an epithelial phenotype (Huels et al., 2015).Thus, we investigated the E-cadherin-β-catenin interaction. CHIR-99021 reduced the proportion of E-cadherin bound β-catenin whilst aspirin treatment promoted the E-cadherin-β-catenin interaction (Fig. 5D). The novel observation that the aspirin-mediated E-cadherin increase is paralleled by greater E-cadherin-β-catenin binding further supports the hypothesis that aspirin promotes an epithelial phenotype by Wnt inhibition.

The addition of CHIR-99021 to organoids grown from FAP non-adenomatous tissue further promoted the cystic phenotype identified in Wnt hyperactive tissues with all organoids appearing cystic after 15 days of treatment. This alteration in cystic: budding organoid ratio was rescued with aspirin (Fig. 5E). Furthermore, aspirin both as a single agent or in combination with CHIR-99021, reduced the expression of β-catenin and the stem cell markers, SOX9 and Lgr5 (Fig. 5F). Taken together, we show that aspirin treatment reverses the Wnt-driven EMT and stem phenotype in both CRC cell lines and organoids derived from human FAP colonic mucosa.

### Aspirin increases expression of the Wnt inhibitor, Dickkopf-1

The loss of endogenous Wnt antagonists during colorectal carcinogenesis is well-recognised and may impact on survival. Dickkopf-1 (DKK-1) is a specific canonical Wnt antagonist and acts as a potent Wnt inhibitor by preventing the LRP6-frizzled interaction and blocking canonical Wnt activation (Semënov et al., 2001). DKK-1 negatively correlates with EMT with DKK-1 overexpression promoting a non-invasive, epithelial phenotype in CRC cells (Qi et al., 2012). Aspirin treatment markedly increases DKK-1 expression whilst decreasing LRP6 expression in CRC cells which is paralleled by a decrease in secreted DKK-1 in the media (Fig. 6A, B). The aspirin-mediated increase in DKK-1 is not attenuated in the presence of CHIR-99021. Aspirin increases nuclear DKK-1 expression in HCT116 cells which is associated with reduced nuclear c-myc and snail protein expression (Fig. 6C). Aspirin treatment increased *DKK-1* transcript expression in organoids derived from FAP non-adenomatous and adenoma tissue (Fig. 6D). Since DKK-1 expression can be silenced due to methylation we used human foetal colonic organoids to minimise potential epigenetic effects in human adult colon. Aspirin treatment of foetal organoids significantly increased *DKK-1* transcript expression (Fig. 6E) and whilst there was no alteration of the cystic phenotype, due to the shorter treatment duration, there was a significant reduction in the average organoid size compared to controls (Fig. 6F). To investigate DKK-1 dependency, DKK-1 was knocked down in HCT116 cells with siRNA (Fig. 6G). The data suggest that the effects on EMT and stem markers are in part due to the aspirin-mediated increase in DKK-1 expression, since siDKK-1 shows an increase in c-myc with a concurrent decrease in E-cadherin expression. Overall, we present data that robustly show that aspirin increases DKK-1 expression in CRC cells and in FAP organoids.

**Figure 6:**
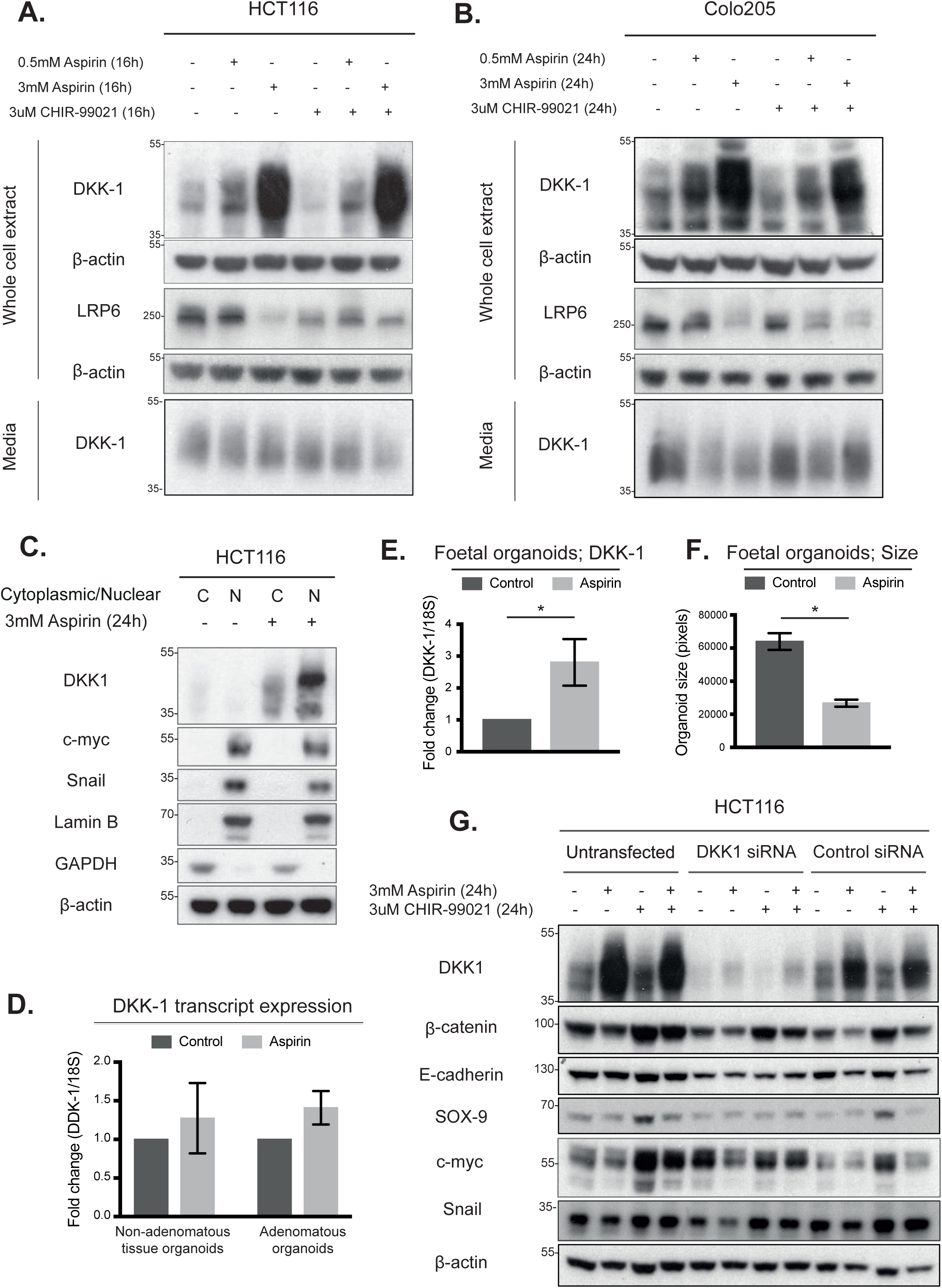
Aspirin treatment increases the expression of intracellular Dickkof-1 whilst reducing secreted Dickkof-1. Immunoblotting for DKK-1 and LRP6 protein from whole cell extracts and secreted DKK-1 protein from media. Cells treated with either 0.5mM aspirin, 3mM aspirin, 3μM CHIR-99021 or combination for 16 and 24 hours in HCT116 (A) and Colo205 (B) cells. Immunoblotting of cytoplasmic/nuclear extracts from HCT116 cells treated with 3mM Aspirin for 24 hours (C). Immunoblotting data representative of three biological replicates. *DDK-1* transcript expression in organoids, derived from FAP non-adenomatous and adenomatous tissue, treated with 2mM Aspirin for 4 hours (D). Combined data from three individual FAP patients. *DDK-1* transcript expression in foetal organoids treated with 2mM Aspirin for 8 days (E). Average size of foetal organoids treated with 2mM Aspirin for 8 days (F). Foetal organoid data represents 4 biological replicates. Immunoblotting of untransfected HCT116 cells and HCT116 cells transfected with either DKK-1 or control siRNA (G) Cells were treated with 3mM aspirin, 3μM CHIR-99021 or combination for 24 hours. B-actin represents sample control. DKK-1 siRNA data representative of three biological replicates. Error bars represent standard error of mean. Significance determined by unpaired students t-test. * p-value <0.05.

## DISCUSSION

Recent evidence has highlighted that aspirin, in addition to its CRC prevention role, may confer a survival benefit following cancer diagnosis. The challenge remains in delineating the signaling pathways responsible and identifying potential markers of response. Here, we show that aspirin rescues the Wnt-driven cystic organoid phenotype and reduces stem cell marker expression in human colon from patients with FAP. Aspirin can reduce the migratory and invasive capabilities of CRC cell lines whilst promoting an epithelial phenotype both *in vitro* and in *in vivo* mouse models. We demonstrate for the first time that aspirin strikingly increases expression of the endogenous Wnt inhibitor, Dickkopf-1 which contributes to the aspirin-mediated inhibition of Wnt signaling, stem-like phenotype inhibition and epithelial phenotype promotion.

Notably, we report the novel observation that aspirin rescues the characteristic aberrant Wnt-driven cystic phenotype by promoting budding in human colonic organoids from patients with FAP. This intestinal organoid phenotype transition has only been observed previously in a mismatch repair deficient mouse model (Keysselt et al., 2017). Increased stem cell, specifically Lgr5, and mesenchymal marker expression is associated with poor survival in CRC (Xu et al., 2018),(Wang et al., 2018). Cell line data suggests that aspirin decreases stem cell expression and function in breast and CRC (Maity et al., 2015; Wang et al., 2017). Here, we establish the *in vivo* relevance using Apc-driven mouse and human FAP organoid models to show that aspirin reduces stem cell marker expression *in vivo*. Paneth cells constitute the niche for Lgr5 stem cells within the small intestine (Sato et al., 2011) and the finding that aspirin decreases the expression of the Paneth cell marker lysozyme in mouse small intestine suggests it may be remodelling the stem cell niche. Paneth cell metaplasia, which has been debated as an early marker of CRC, was less frequently observed in colonic mucosa of aspirin users (Zagorowicz et al., 2013). The novel findings of aspirin-induced cystic phenotype rescue and reduction in stem cell marker expression in human tissue are highly relevant, given the critical role of hyper-activated Wnt signaling in CRC, and highlight their potential role as phenotypic biomarkers of response.

In addition to regulating the stem cell niche, Wnt signaling drives EMT in collaboration with key pathways including TGF-β and mTOR. Moreover, highlighting a feedforward loop, EMT and CSC induction in breast tissue increased Wnt activity and reduced DKK-1 expression (Scheel et al., 2011). The ability of aspirin to decrease cellular migration and invasion whilst promoting an epithelial phenotype is consistent with previous literature. Aspirin reduced lung cancer cell migration by E-cadherin upregulation via slug repression (Khan et al., 2016). Aspirin also decreased pro-invasive metalloproteinase expression in CRC cells (Koontongkaew et al., 2010). Recently, it was reported that supra-physiological aspirin reduced CRC cell migration and attenuated platelet-mediated EMT (Ying et al., 2018),(Guillem-Llobat et al., 2016). Importantly, we show that aspirin-induced inhibition of migration, invasion and motility and the promotion of an epithelial phenotype occurs at physiologically relevant doses independently of platelet inhibition in vitro. The role of platelets in EMT is biologically relevant given that aspirin is an anti-thrombotic agent and platelet inhibition may contribute to the effects of aspirin in the Apc mouse models.

We demonstrate that Wnt activation, using the GSK-3β inhibitor, induces EMT and stem-like alterations in CRC cells, which are rescued by aspirin. Our results show that aspirin increases expression of the endogenous secreted Wnt antagonist and β-catenin transcription target, DKK-1(González-Sancho et al., 2005). This negative feedback is lost during CRC due to DKK-1 promoter hypermethylation and demethylation restores DKK-1 expression and tumour suppressing function (Aguilera et al., 2006). Its role in intestinal homeostasis is further highlighted by knock-down leading to epithelial cell hyperproliferation in murine intestine(KOCH et al., 2011). DKK-1 over-expression reduces cellular migration and invasion whilst promoting an epithelial phenotype and reduces stem cell marker expression, including Lgr5, in CRC cells (Qi et al., 2012). In humans, high serum DKK-1 correlates with increasing CRC stage (Gurluler et al., 2014) whereas tissue DKK-1 expression is lost with cancer progression. Here, we demonstrate that aspirin robustly increases DKK-1 tissue expression and decreases secreted DKK-1 levels in CRC models. Thus, raising the possibility of using serum DKK-1 as a biomarker of response to aspirin. Interestingly, aspirin reduces plasma DKK-1 levels in non-cancer diabetic patients (Lattanzio et al., 2014) and exercise also reduces DKK-1 serum levels in breast cancer survivors (Kim et al., 2017).

Several potential mechanisms underlying the aspirin-mediated increase in DKK-1 remain to be explored. Aspirin has been shown to reduce methylation of promoter-associated CpGs in normal colon mucosa (Noreen et al., 2014). LSD1 is a demethylase that is highly expressed in CRC compared to normal colon with knockdown leading to decreased tumorigenic properties and increased DKK-1 expression (Huang et al., 2013). It is possible that aspirin increases DKK-1 expression through LSD1 inhibition which demethylates DKK-1. However, it may also suggest that aspirin targets the Wnt signaling pathway at multiple levels, which could be a therapeutic benefit. Indeed this has been demonstrated with aspirin reported to inhibit EP300, a transcriptional co-activator of β-catenin (Pietrocola et al., 2018). Using mouse and human CRC models our data identify novel phenotypic biomarkers of response to aspirin with an increased epithelial and reduced stem-like state potentially mediated by an increase in the Wnt antagonist DKK-1. These observations reveal a novel mechanism of aspirin-mediated Wnt inhibition and potential biomarkers for chemoprevention and adjuvant aspirin human trials.

## MATERIALS AND METHODS

### Cell Culture

Colorectal cancer cell lines (HCT116 and Colo205) are available from the American type culture collection (ATCC). Cells were maintained in either McCoy’s (HCT116) or DMEM (Colo205) media supplemented with 10% foetal calf serum, 100IU/ml penicillin and 100μg/ml streptomycin.

### Wound healing assays

A wound was created in a confulent cell monolayer using a p200 pipette tip. Fresh media (containing either 0.5% or 10% serum) and treatment was added with images taken at 0, 24 and 48 hours using a Zeiss Axiovert 100 microscope with a 4x objective.

### Single cell motility assays

Cells were seeded sub-confulently in medium containing either 0.5% or 10% serum. Treatment was added and images taken on a live cell imaging microscope, Zeiss Axiovert 200. A 10x objective was used with images captured every 30 minutes for 24 hours. Cell movement was measured and calculated using a manual tracking and chemotaxis plugins for Image J.

### Organotypic invasion assays

Organotypic collagen based invasion assays were used as previously described (Timpson et al., 2011). Briefly, cells invaded collagen-fibroblast matrices for 7 days. These matrices were fixed in 4% paraformaldehyde, paraffin embedded and sections were stained with haematoxylin and eosin. Cells were imaged using a Zeiss Axioplan 2 microscope with a 10x objective and couted using Image J software. Telomerase immortalised fibroblasts were kindly provided by John Dawson (IGMM, Edinburgh).

### Immunoblotting

Cells were lysed in whole cell lysis buffer (Sup. Table 1) for 45 minutes and clarified at 13000rpm for 20mins. For cytoplasmic and nuclear fractioned extractions, cell pellets were lysed in cytoplasmic lysis buffer for 10 minutes then centrifuged at 3000rpm for 10mins. The nuclear pellet was lysed in hypotonic buffer for 30 minutes then clarified at 13000rpm for 20mins. Protein extractions were quantified by Bradford’s method (Bio-Rad). Protein lysates were seperated by SDS-PAGE and transferred to PDVF membranes before blocking with 5% non-fat milk. Primary antibodies were incubated for 16hours at 4°C before secondary HRP-conjugated antibodies were incubated for 1 hour (antibodies are detailed in Sup. Table 2). Antigen-antibody complexes were visualised using chemilumunescence (Santa-cruz).

### Immunoprecipitation

Cells were lysed in immunoprecipitation lysis buffer (Sup. Table 3) for 45 minutes and clarified at 13000rpm for 20mins. For each reaction, 5μl of either E-cadherin or β-catenin antibody (BD Transduction) was added to 800μg of protein and incubated at 4°C for 24 hours. Dynabeads protein G (50μl) were added to each reaction and incubated for 16 hours at 4°C. Beads were washed in PBS with 0.02% Tween-20 three times then eluted in 100μl of SDS sample buffer at 70°C for 10mins. Samples were immunoblotted on SDS-PAGE.

### Pull-down of secreted DKK-1

DKK-1 protein was pulled down from cell media by StrataClean Resin (Agilent Technologies) in accordance with manufacturers’ instructions and detected by immunoblotting using anti-DKK-1 antibody.

### Immunofluorescence

Cells were seeded on glass coverslips. Following treatment, cells were fixed in 4% paraformaldehyde for 15 minutes, incubated in permeabilisation buffer (0.2% Triton X) for 10 minutes then blocking buffer (3% goat serum, 3% bovine serum albumin) for 1 hour. E-cadherin or Zona-occludens 1 antibody (CST), diluted 1:200, was incubated overnight at 4°C before incubation with Alexafluor488 rabbit antibody (Life technologies) at 1:1000 for 1 hour. For F-actin staining, after blocking, cells were incubated with AlexaFlur488 phalloidin (Life technologies) for 1 hour at room temperature then mounted using Vectashield mouting medium with DAPI (Vector laboratories). Cells were imaged on Zeiss Axioplan 2 using 10x, 40x and 100x objectives. Mean grey area of image was divided by number of nuclei per image to determine a value for staining intensity.

### Quantitative real time polymerase chain reaction (qRT-PCR)

Following treatment, RNA was extracted using Ribopure RNA isolation kit (Life technologies). RNA was DNase treated before cDNA was synthesised using M-MLVRT (Promega). Quantitative real time PCR was completed on Lightcycler 480 (Roche) using SYBR green master mix (Life technologies). Primer sequences can be found in Sup. Table 4.

### Organoid culture

The culture of colonic organoids from human tissue has been described previously (Sato et al., 2011). Briefly, human normal colon mucosa was collected during tumour resection, crypts disocciated and seeded in matrigel. Experimental methodology is detailed in supplementary.

### Animal studies

All animal experiments were approved by either the University of Edinburgh or University of Glasgow ethics committee and preformed under a UK Home Office project licence. Three Apc^Min/+^ mouse cohorts were used; a 4 week treatment group, a 21 day treatment group and a 7 day treatment group. Experiments detailed in suplementary.

### Immunohistochemistry

Sections were deparaffinised and rehydrated by repeated washes in xylene and ethanol. Antigen retrieval was completed by incubation in boiling 10% citrate buffer (Thermofisher). Endogenous blocking was achieved with incubation in 3% hydrogen peroxide (Sigma-Aldrich) for 10 minutes followed by 5% goat serum (Thermofisher) for 1 hour. Primary antibody was incubated for 16hours at 4°C before secondary rabbit antibody (Dako) added for 30 minutes. Signal was detected using DAB peroxidase substrate kit (Vector laboratories). Sections were counterstained with Haematoxylin and mouted using DPX mounting medium (Sigma-Aldrich). All slides were imaged using a Hamamatsu nanozoomer. Staining intensity was graded as 0-negative, 1-mild, 2-moderate, 3-intense and multiplied by percentage of adenoma expressing that intensity to give arbitury values per adenoma.

### RNAscope

RNAscope is commercially available from Advanced Cell Diagnostics (ACD). Here, the probes used were: Mouse TROY (#420241) and Mouse Lgr5 (#312171). RNAscope staining was performed according to manufacturer’s instructions. Quantification of RNAscope from adenoma tissue was completed manually using Image J software with all dots within an adenoma counted and normalised to the area. Quantification of RNAscope from organoids was completed using Qupath software as previously described ([CSL STYLE ERROR: reference with no printed form.]). All well-formed organoids present on the slide were analysed by setting threshold values (to account for the differential DAB staining intensities) and then calculating the number of positive cells.

### Transfection of siRNAs

Cells were transfected with 25 nM siRNA that targets human *DKK-1* (Cat number E-003843-00-0005, Dharmacon) or nonspecific siRNA (Cat number E-001910-01-5, Dharmacon) as a negative control using Lipofectamine2000 (Invitrogen by Life Technologies) in accordance with the manufacturers’ instructions. Then, the cells were cultured for additional 24 hours, and treated with aspirin and/or CHIR-99021 for 24 hours.

## Abbreviations

APC: Adenomatous polyposis coli
CRC: Colorectal cancer
CSC: Cancer stem cells
DKK-1: Dickkopf-1
EMT: Epithelial-Mesenchymal transition
F-actin: Filamentous actin
FAP: Familial adenomatous polyposis
GSK-3B: Glycogen synthase kinase 3B
LEF: Lymphoid enhancer factor
Lgr5: Low-density lipoprotein receptor-related protein 6
LRP6: Leucine-rich repeat-containing G-protein coupled receptor 5
mTOR: Mechanistic target of rapamycin
NSAID: Non-steroidal anti-inflammatory drug
PBS: Phosphate buffered saline
PDVF: Polyvinylidene difluoride
Rac1: Ras-related C3 botulinum toxin substrate 1
RhoA: Ras homolog gene family, member A
RNA: Ribonucleic acid
ROCK1: Rho-associated protein kinase 1
ROCK2: Rho-associated protein kinase 2
SDS-PAGE: Sodium dodecyl sulphate polyacrylamide gel electrophoresis
TCF: T-cell factor
TGF-B: Transforming growth factor beta
TROY: Tumour necrosis factor receptor superfamily member 19
ZO-1: Zona-occludens 1

## Author Contributions

Study concept and design (FD). Acquisition of data (KD, AV, PFV, VR, AL, AMOF). Analysis and interpretation of data (KD, AV, KM and FD). Drafting of manuscript (KD and FD). Critical revision of the manuscript for important intellectual content (KD, AV, OS, KM, SMF, MJA, MGD and FD). Technical support (TJ, AMOF, VR, JB, MJA) and material support (KM).

---

**Video 1a:** Control Apc^flox/flox^ organoid over a 48hour period with images captured every hour with a 10x objective. Demonstrates a cystic organoid which grows but doesn’t exhibit any bud formation.

**Video 1b:** Aspirin treated Apc^flox/flox^ organoid over a 48hour period with images captured every hour with a 10x objective. Organoids treated with 2mM Aspirin with images captured on day 2 and 3 of treatment. Demonstrates organoids exhibiting the initial stages of bud formation.

